# Test-Retest Reproducibility of Single- and Cross-Population White Matter Atlases in Diffusion MRI Tractography

**DOI:** 10.64898/2026.09.11.750840

**Authors:** Yijie Li, Xiaofan Wang, Jincheng Sun, Wei Zhang, Ye Wu, Li Yin, Yuqian Chen, Yogesh Rathi, Nikos Makris, Lauren J. O’Donnell, Fan Zhang

## Abstract

Diffusion MRI tractography enables noninvasive mapping of white matter fiber tracts. Atlas-based white matter parcellation supports automated tract identification by assigning individual streamlines to atlas-defined clusters and anatomical tract labels. Because clustering-based white matter atlases are constructed from cohort-specific tractography data, they capture common white matter organization represented in the atlas-construction population. Although major white matter anatomy is shared across populations, subtle population-related anatomical variability may influence atlas representation and test-retest correspondence. Therefore, the reproducibility and cross-population generalizability of tractography-based white matter atlases are important considerations for quantitative neuroimaging studies. In this study, we evaluated whether incorporating cross-population anatomical variability during atlas construction improves test-retest reproducibility. To do so, we compared a single-population ORG atlas constructed from a Western cohort with the cross-population East-West White Matter Atlas constructed from both Eastern and Western cohorts. Test-retest diffusion MRI scans from the Human Connectome Project Young Adult (HCP-YA) dataset and the Connectivity-based Brain Imaging Research Database (C-BIRD) were analyzed as independent Western and Eastern test-retest cohorts, respectively. Whole-brain tractography was reconstructed for each dMRI scan and parcellated using both atlases. Reproducibility was assessed using tract detection rate at both cluster level and anatomical tract level, weighted Dice coefficient, and the relative difference of mean fractional anisotropy (FA). Both atlases showed stable tract detection across test-retest scans in both cohorts. Compared with the ORG atlas, the East-West White Matter Atlas achieved higher overall spatial overlap and lower test-retest variability in mean FA, although atlas performance varied across individual tracts. These findings suggest that integrating cross-population information during atlas construction improves the reproducibility and generalizability of white matter atlas mapping across independent populations and imaging protocols.

## 1 Introduction

Diffusion magnetic resonance imaging (dMRI) tractography has become an essential technique for investigating the structural connectivity of the human brain in vivo (Basser et al., 1994). By estimating white matter orientations from diffusion signals and reconstructing their trajectories, tractography enables noninvasive mapping of brain connectivity and provides a powerful framework for characterizing the organization of white matter architecture (Basser et al., 2000; Zhang et al., 2022). Over the past two decades, dMRI tractography has been widely applied in studies of brain development, aging, neurological and psychiatric disorders, and neurosurgical planning (Piper et al., 2014; Pannek et al., 2014; Essayed et al., 2017; Zhang et al., 2026). Despite these advances, whole-brain tractography typically produces a large number of streamlines with complex spatial configurations, making direct interpretation, anatomical labeling, and quantitative comparison across subjects challenging (Maier-Hein et al., 2017; Thomas et al., 2014; Poulin et al., 2019). These challenges have motivated the development of automated and standardized approaches for organizing tractography data into anatomically meaningful white matter structures (O’Donnell et al., 2013; Garyfallidis et al., 2018; Wassermann et al., 2016).

Fiber clustering-based white matter atlases provide an effective framework for the automated organization and quantitative analysis of whole-brain tractography data (O’Donnell and Westin, 2007; Chen et al., 2023; Guevara et al., 2011; Garyfallidis et al., 2012). These atlases group streamlines into anatomically meaningful fiber bundles according to their geometric similarity and spatial organization, thereby enabling automated white matter parcellation and standardized tract-specific measurements across subjects and studies (Zhang et al., 2018; Guevara et al., 2012). Compared with approaches that segment a predefined set of tracts using specified anatomical regions or rules, clustering-based atlases enable a more fine-grained and standardized organization of whole-brain tractography (Zhang et al., 2019). Importantly, most tractography atlases are population-oriented because they are constructed from tractography data acquired from specific cohorts to capture common patterns of white matter organization (Zhang et al., 2018). Therefore, the anatomical variability represented by an atlas may be influenced by the population used for atlas construction, as well as by differences in sample composition, imaging protocols, clustering procedures, and anatomical labeling strategies (Yang et al., 2020a,b). Atlases derived from relatively homogeneous or single-population datasets may provide accurate representations for cohorts similar to the atlas population, whereas atlases constructed from multiple populations may better capture inter-population anatomical variability and improve generalizability (Zhang et al., 2026; Li et al., 2024).

Population diversity has been increasingly recognized as an important factor in brain atlas construction. Studies of brain template construction and cortical morphology have shown that population-related anatomical variation can affect atlas representation and anatomical correspondence, suggesting that atlases derived from a single population may not always generalize optimally to independent cohorts (Liang et al., 2015; Yang et al., 2020a,b). This consideration is also relevant to dMRI tractography-based white matter atlases, where an atlas constructed from a single population may provide less comprehensive mapping when applied to independent populations. In contrast, cross-population white matter atlases integrate tractography information from multiple populations to capture both shared fiber organization and inter-population anatomical variability, potentially providing a more robust anatomical reference for concurrent white matter mapping across different cohorts (Li et al., 2024; Zhang et al., 2026). However, whether cross-population atlas construction improves test-retest reproducibility compared with single-population atlases across independent cohorts remains to be systematically evaluated.

Reliability and reproducibility are critical considerations for diffusion MRI tractography, particularly when tractography-derived measurements are used for quantitative neuroimaging analyses. Tractography results can be influenced by multiple sources of variability, including image acquisition protocols, scanner differences, diffusion reconstruction methods, tractography parameters, registration procedures, and parcellation strategies. Test-retest reproducibility has therefore become an important criterion for evaluating the stability and robustness of tractography-derived measurements (Boukadi et al., 2019). In the context of atlas-based white matter mapping, reproducibility refers to the ability to consistently identify corresponding white matter structures and obtain stable geometric and diffusion-derived measurements across test-retest scans of the same subject (Zhang et al., 2019). In this study, we investigated whether cross-population white matter atlasing improves test-retest reproducibility compared with single-population atlas construction. Specifically, we used the ORG atlas and the East-West White Matter Atlas as representative examples of two atlas construction strategies: a single-population atlas derived from a Western cohort (Zhang et al., 2018) and a cross-population atlas constructed by integrating tractography data from both Eastern and Western cohorts (Li et al., 2024). The Connectivity-based Brain Imaging Research Database (C-BIRD) from the Beijing Normal University dataset (Lin et al., 2015) and the Human Connectome Project Young Adult (HCP-YA) dataset (Van Essen et al., 2013) were analyzed as independent Eastern and Western test-retest cohorts, respectively. By evaluating tract detection, spatial overlap, and diffusion-derived measurement stability across these cohorts, this study aims to investigate whether incorporating cross-population information during white matter atlas construction enhances the robustness, reproducibility, and generalizability of quantitative tractography analyses.

## 2 Methods

### 2.1 Datasets, data processing, and tractography

We evaluated test-retest reproducibility using the two public dMRI datasets described above: HCPYA and C-BIRD. Together, the study included 101 young adult participants aged 19-35 years, comprising 61 females and 40 males. The two datasets were acquired using different diffusion MRI protocols, allowing atlas reproducibility to be assessed across independent cohorts and imaging conditions. Detailed demographic information, diffusion acquisition parameters, and test-retest intervals are summarized in Table 1.

**Table 1:** Demographic information and dMRI acquisition parameters of the datasets included in this study.

| Dataset | Subjects | Age | Gender | dMRI acquisition parameters | Test-retest interval |
| --- | --- | --- | --- | --- | --- |
| HCP-YA | 44 | 22-35 years;<br>$30.4 \pm 3.3$<br>years | 31 F; 13 M | 18 b0 images; 90 directions; $b = 3000 \text{ s/mm}^2$; TE/TR = 89/5520 ms; 1.25 mm isotropic | 18-328 days;<br>$134 \pm 62$ days |
| C-BIRD | 57 | 19-30 years;<br>$23.1 \pm 2.3$<br>years | 30 F; 27 M | 1 b0 image; 30 directions; $b = 1000 \text{ s/mm}^2$; TE/TR = 89/8000 ms; 2.2 mm isotropic | 33-55 days; $41 \pm 5$ days |

For each subject, both test and retest dMRI scans were preprocessed to reduce potential artifacts. Standard dMRI preprocessing procedures included brain masking, correction for eddy current-induced distortions, motion correction, and EPI distortion correction. For the HCP-YA dataset, we used dMRI data that had already been processed with the HCP minimal preprocessing pipeline (Glasser et al., 2013). For the C-BIRD dataset, motion and eddy current-induced distortion corrections were performed using DTIPrep v1.2.11 (https://www.nitrc.org/projects/dtiprep) (Oguz et al., 2014). Volumes with translational displacement greater than 2 mm or rotational displacement greater than 2 degrees were automatically identified and excluded from further analysis. Brain extraction was then performed using CNN-Diffusion-MRIBrain-Segmentation v0.3 (https://github.com/pnlbwh/CNN-Diffusion-MRIBrain-Segmentation/tree/v0.3) (Cetin-Karayumak et al., 2024).

Whole-brain tractography was performed using the two-tensor unscented Kalman filter (UKF) tractography method, as implemented in the ukftractography package (https://github.com/pnlbwh/ukftractography) (Malcolm et al., 2010; Chen et al., 2015). The UKF method fits a two-tensor mixture model to the dMRI data during fiber tracking and incorporates prior information from the previous tracking step to stabilize model estimation. UKF tractography was selected because it has demonstrated high consistency in reconstructing white matter streamlines across independently acquired populations with varying ages, health conditions, and imaging protocols (Chen et al., 2016; Zhang et al., 2026). For each subject, approximately 1,000,000 streamlines were generated from each dMRI scan using the default parameters.

For each subject, tractography datasets generated from the test and retest scans were aligned into the same space. Specifically, the fractional anisotropy (FA) image derived from the retest scan was registered to the FA image derived from the test scan using the Advanced Normalization Tools (ANTs) package (https://github.com/ANTsX/ANTs) (Avants et al., 2009). FA-based registration was used because FA images are sensitive to white matter architecture and can provide reliable anatomical correspondence between diffusion MRI datasets (Goodlett et al., 2006; Besseling et al., 2012). The resulting transformations were applied to the tractography data generated from the retest scan in 3D Slicer, thereby transforming the retest tractography into the test-scan space (Norton et al., 2017; Zhang et al., 2020). Because the transformations were applied after tractography reconstruction, this procedure avoided resampling or blurring of the original diffusion-weighted image data. An overview of the complete analysis workflow is shown in Figure 1.

**Figure 1.**
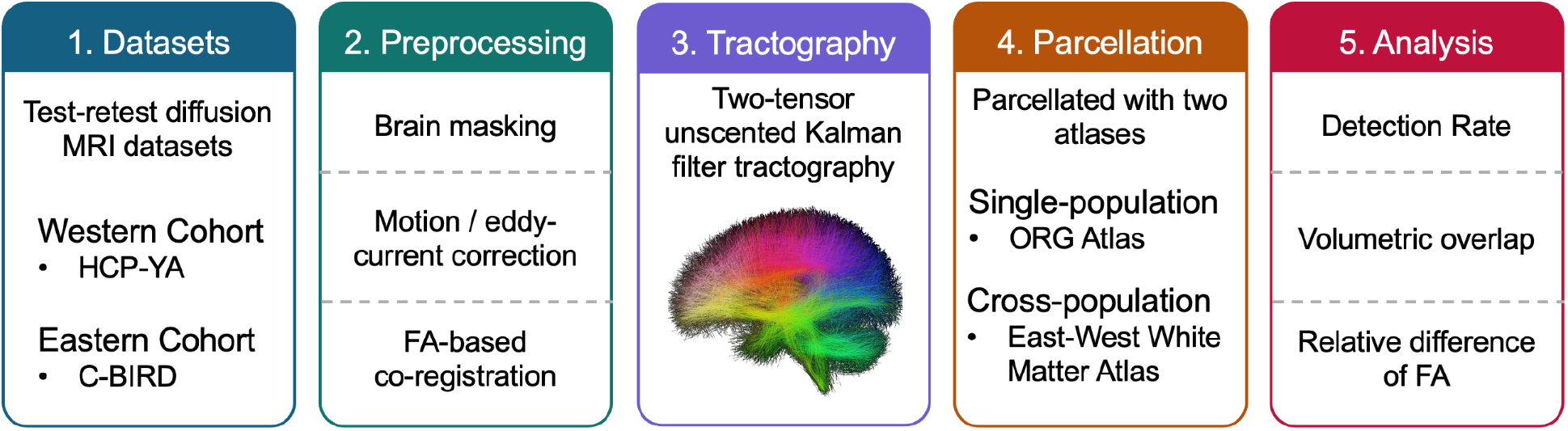
Study workflow for test-retest reproducibility analysis.

### 2.2 White Matter Parcellation

After whole-brain tractography was obtained, white matter parcellation was performed using a fiber clustering approach. For each subject, individual whole-brain tractography data were parcellated using two clustering-based white matter atlases: the ORG atlas (Zhang et al., 2018) and the East-West White Matter Atlas (Li et al., 2024). Both atlases provide an 800-cluster whole-brain white matter parcellation and anatomically annotated fiber-tract labels, but they differ in the populations used for atlas construction.

The ORG atlas was constructed from dense tractography maps of 100 HCP-YA subjects by grouping fibers across subjects according to similarities in shape and spatial location. It includes anatomical labels for 58 deep white matter fiber tracts and 198 short- and medium-range superficial fiber tract groups, with superficial tracts organized into 16 categories according to the cortical lobes they connect. In contrast, the East-West White Matter Atlas was constructed from a cross-population cohort of 306 subjects from the Chinese Human Connectome Project (CHCP) and HCP-YA datasets (Ge et al., 2023; Van Essen et al., 2013), including 153 subjects from each population. It provides anatomical labels for 60 deep white matter tracts and 222 superficial fiber tract groups, with superficial tracts similarly organized into 16 lobe-based categories. In both atlases, anatomical labels were assigned based on expert neuroanatomical knowledge. Therefore, the ORG atlas was used as a representative single-population atlas derived from a Western cohort, whereas the East-West White Matter Atlas was used as a representative cross-population atlas integrating Eastern and Western cohorts.

For each subject, atlas-based fiber clustering was used to assign streamlines from individual tractography data to the corresponding atlas-defined clusters (O’Donnell et al., 2012; O’Donnell and Westin, 2007; Zhang et al., 2018). Briefly, tractography-based registration was first performed to align the subject’s tractography data to the atlas space. Fiber spectral embedding was then applied to compute fiber similarity between the subject and the atlas, after which each subject streamline was assigned to its most corresponding atlas cluster. Anatomically defined white matter tracts were then obtained by grouping atlas clusters according to the tract annotations provided by each atlas. For the ORG atlas, the 800 atlas clusters consisted of 84 commissural clusters and 716 hemispheric clusters. Because hemispheric clusters were represented separately in the left and right hemispheres, this yielded 1,516 hemisphere-separated cluster units. For the East-West White Matter Atlas, the 800 clusters consisted of 99 commissural clusters and 701 hemispheric clusters, yielding 1,501 hemisphere-separated cluster units. All fiber clustering and white matter parcellation procedures were performed using the whitematteranalysis software package (https://github.com/SlicerDMRI/whitematteranalysis), with all parameters set to their default values.

### 2.3 Test-retest measurements

After white matter parcellation, test-retest measurements were computed for the parcellated white matter structures to evaluate reproducibility. Three complementary aspects of reproducibility were assessed: detection reproducibility, geometric reproducibility, and diffusion-derived microstructural measurement reproducibility.

For detection reproducibility, the detection rate was quantified at two levels. First, the cluster-level detection rate was calculated using hemisphere-separated atlas cluster units, in which left- and right-hemisphere clusters were counted separately when applicable. This analysis evaluated the stability of fine-grained atlas cluster detection across test-retest scans. Second, the anatomical tract-level detection rate was calculated using anatomically defined white matter tracts. This analysis evaluated whether major anatomical fiber tracts could be consistently identified across test-retest scans. At both levels, a structure was considered detected if it contained at least 10 streamlines (Chen et al., 2023; Zhang et al., 2018), and the detection rate was computed as the proportion of detected structures across subjects for each atlas and scan.

For geometric reproducibility, we used the weighted Dice coefficient (wDice) to quantify the spatial overlap between corresponding parcellated white matter tracts from the test and retest scans (Dice, 1945; Cousineau et al., 2017; Zhang et al., 2019). Unlike the standard Dice coefficient, wDice incorporates voxel-wise fiber density weights and therefore gives a greater contribution to voxels traversed by denser fiber populations. A higher wDice value indicates greater spatial reproducibility between corresponding parcellated white matter structures.

For diffusion-derived microstructural measurement reproducibility, we evaluated the reproducibil-ity of mean FA within each parcellated white matter structure (Papinutto et al., 2013; Zhang et al., 2019). Specifically, mean FA was computed from the voxels traversed by the fibers of each structure. Following strategies used in related work, test-retest variability was quantified using a normalized relative difference (RD), calculated as the absolute difference between the test and retest mean FA values divided by their sum. A lower RD value indicates higher reproducibility. This normalized measure was used because diffusion properties such as FA vary across white matter structures; normalizing the absolute difference by the sum of the two mean values provides a scale-adjusted measure that enables more comparable reproducibility estimates across structures.

### 2.4 Statistical analysis

Statistical comparisons between the two white matter atlases were performed based on the computed test-retest measurements. All statistical analyses were conducted separately for each dataset.

For detection reproducibility, statistical analyses were performed using the detection rates calculated at the cluster and anatomical tract levels, as described above. For each atlas, paired comparisons between the test and retest sessions were performed to assess whether the detection rate differed across test-retest scans.

For anatomical tract-level analyses, we focused on the common deep white matter tract set defined by both atlases and summarized in Table 2. Statistical comparisons were performed using two complementary approaches: an overall tract-level comparison across anatomical tracts and an individual-tract comparison. For the overall comparison, each test-retest measurement was first averaged across subjects for each tract and each atlas. The resulting tract-wise mean values were then compared between the two atlases using two-tailed paired Student’s t-tests, with tracts treated as paired observations. This analysis assessed whether one atlas showed higher overall reproducibility across the tract set. For the individual-tract comparison, each tract was analyzed separately using subject-level test-retest measurements. For each tract, wDice and relative difference values from the same subjects were compared between atlases using two-tailed paired Student’s t-tests. The false discovery rate (FDR) procedure was applied to correct for multiple comparisons across tracts.

**Table 2:** Anatomical tracts included in tract-level reproducibility analyses.^a/sup>^.

| Tract category | Tract names |
| --- | --- |
| Association tracts<br>(22) | Arcuate fasciculus (AF) - L/R; cingulum bundle (CB) - L/R; external capsule (EC) - L/R; extreme capsule (EmC) - L/R; inferior longitudinal fasciculus (ILF) - L/R; inferior occipito-frontal fasciculus (IoFF) - L/R; middle longitudinal fasciculus (MdLF) - L/R; superior longitudinal fasciculus I (SLF I) - L/R; superior longitudinal fasciculus II (SLF II) - L/R; superior longitudinal fasciculus III (SLF III) - L/R; uncinate fasciculus (UF) - L/R |
| Commissural tracts<br>(7) | Corpus callosum 1 (CC 1) - C; corpus callosum 2 (CC 2) - C; corpus callosum 3 (CC 3) - C; corpus callosum 4 (CC 4) - C; corpus callosum 5 (CC 5) - C; corpus callosum 6 (CC 6) - C; corpus callosum 7 (CC 7) - C |
| Projection tracts<br>(20) | Corticospinal tract (CST) - L/R; corona radiata-frontal (CR-F) - L/R; corona radiata-parietal (CR-P) - L/R; striato-frontal (SF) - L/R; striato-occipital (SO) - L/R; striato-parietal (SP) - L/R; thalamo-frontal (TF) - L/R; thalamo-occipital (TO) - L/R; thalamo-parietal (TP) - L/R; thalamo-temporal (TT) - L/R |
| Cerebellar tracts<br>(11) | Cortical-ponto-cerebellar tract (CPC) - L/R; intracerebellar input and Purkinje tract (CBLM-I&P) - L/R; intracerebellar parallel tract (CBLM-PaT) - L/R; middle cerebellar peduncle tract (MCP) - C; inferior cerebellar peduncle tract (ICP) - L/R; superior cerebellar peduncle tract (SCP) - L/R |

<sup>a</sup> A total of 60 anatomical tracts were analyzed, including 22 hemisphere-separated association tracts, 7 commissural tracts, 20 hemisphere-separated projection tracts, and 11 cerebellar tracts. For hemisphere-separated tracts, L/R indicates that the left and right hemispheres were analyzed separately; C indicates commissural tracts.

## 3 Results

### 3.1 Detection rate results

Figure 2 shows the cluster-level detection rates in the test and retest scans for the C-BIRD and HCP-YA datasets. In the C-BIRD dataset, the East-West White Matter Atlas showed higher cluster-level detection rates than the ORG atlas in both test and retest scans. In the HCP-YA dataset, the East-West White Matter Atlas showed slightly lower cluster-level detection rates than the ORG atlas; however, detection rates remained high for both atlases, about 95% in both testretest scans. For both datasets and both atlases, paired t-tests showed no significant differences in cluster-level detection rate between the test and retest scans, indicating stable detection across test-retest acquisitions.

**Figure 2.**
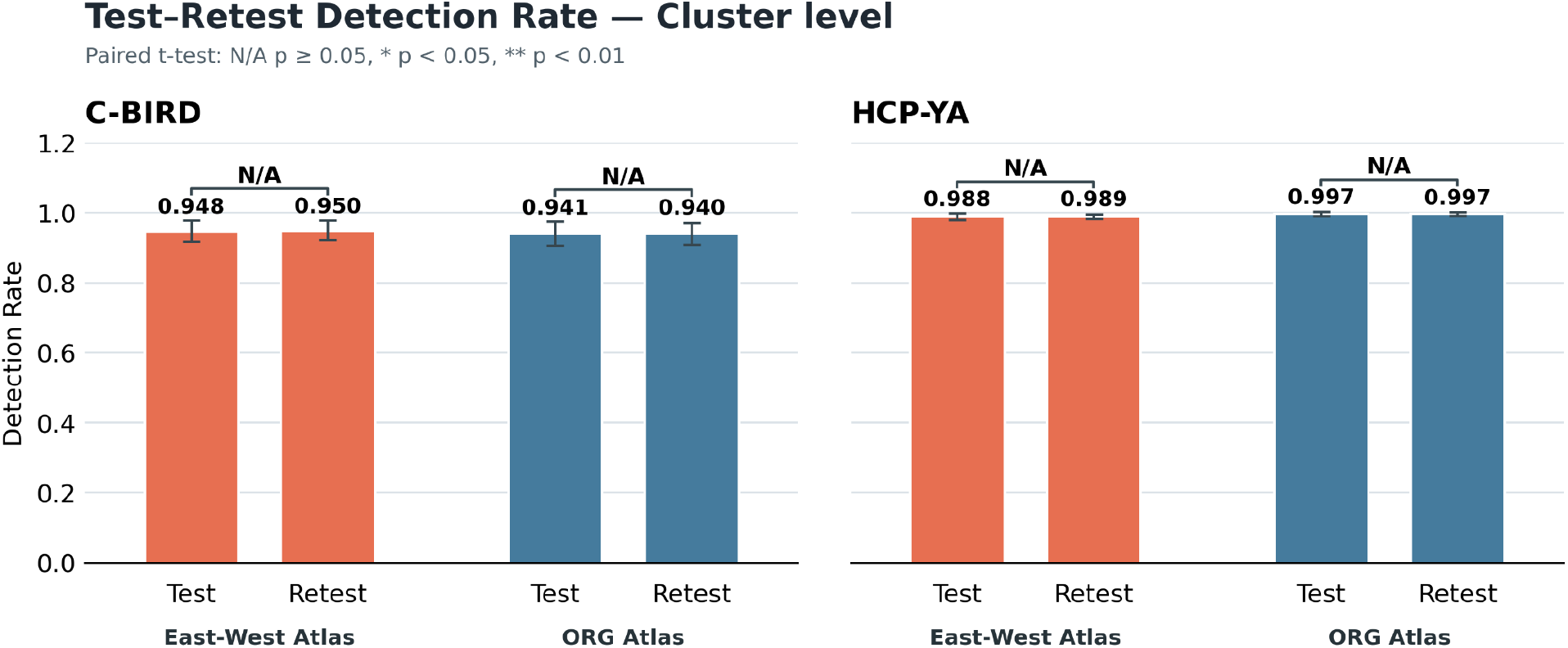
Cluster-level detection rates in the test and retest scans.

Figure 3 shows the anatomical tract-level detection rates. Both atlases achieved complete anatomical tract-level detection in the test and retest scans of both datasets. No significant differences in anatomical tract-level detection rate were observed between test and retest scans for either atlas in either dataset, indicating high test-retest stability of anatomical tract detection.

**Figure 3.**
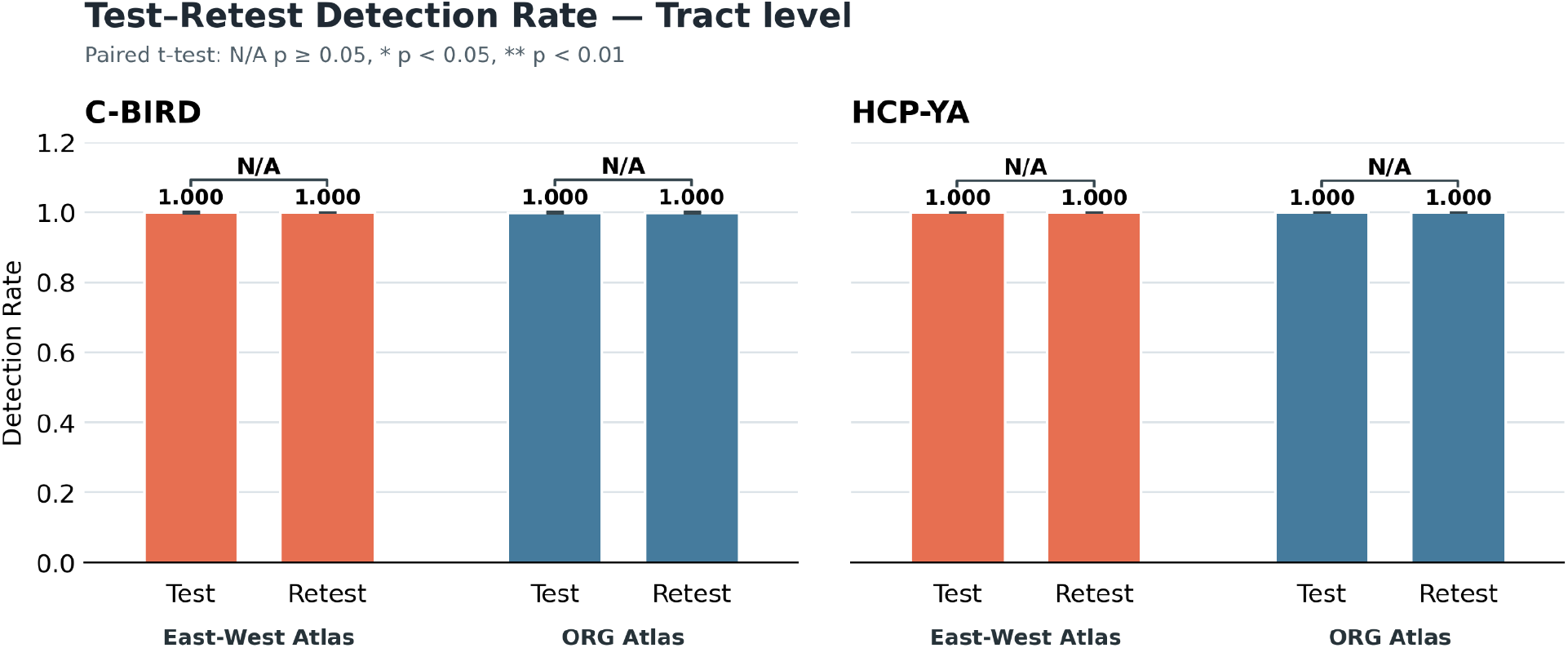
Anatomical tract-level detection rates in the test and retest scans.

### 3.2 Volumetric overlap results

The overall volumetric overlap comparison is shown in Figure 4. In both the C-BIRD and HCP-YA datasets, the East-West White Matter Atlas achieved mean wDice scores above 0.93, indicating high spatial overlap between test and retest parcellations. Compared with the ORG atlas, the East-West White Matter Atlas showed higher overall wDice scores in both datasets. These differences were statistically significant in the overall tract-level comparison (p *<* 0.001, two-tailed paired t-test), indicating improved spatial reproducibility across test-retest scans.

**Figure 4.**
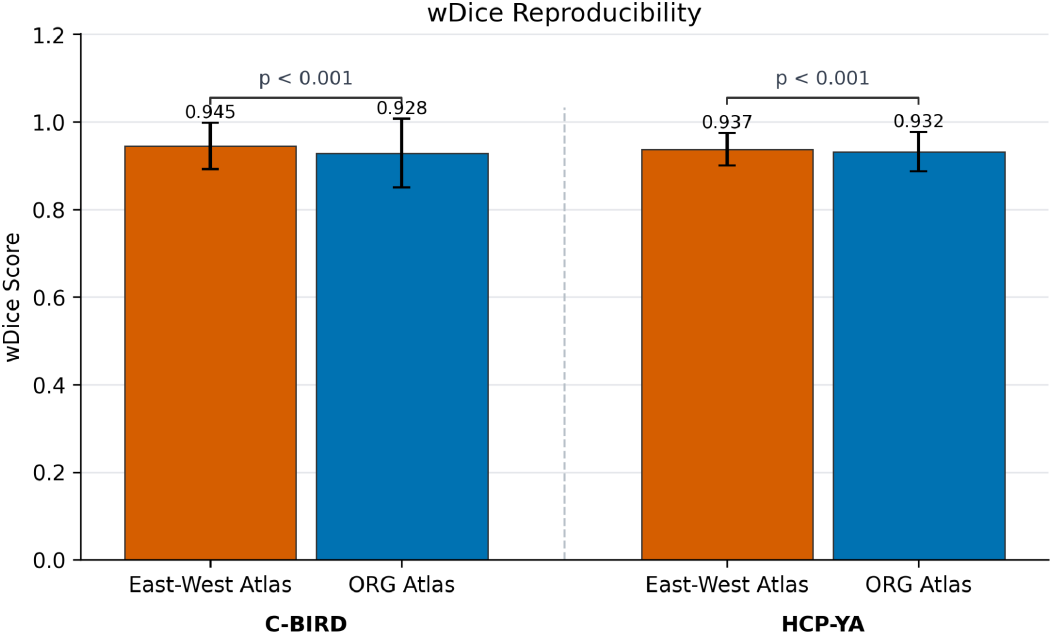
Overall volumetric overlap comparison.

In the individual-tract comparison shown in Figure 5, 39 and 21 tracts in the C-BIRD and HCP-YA datasets, respectively, showed significantly higher wDice scores when parcellated using the East-West White Matter Atlas. In contrast, 6 and 20 tracts in the C-BIRD and HCP-YA datasets, respectively, showed significantly higher wDice scores when parcellated using the ORG atlas (p *<* 0.05, two-tailed paired t-test, FDR-corrected). These findings indicate that the East-West White Matter Atlas generally improved volumetric overlap reproducibility across common deep white matter tracts, although the ORG atlas showed higher reproducibility in a subset of individual tracts.

**Figure 5.**
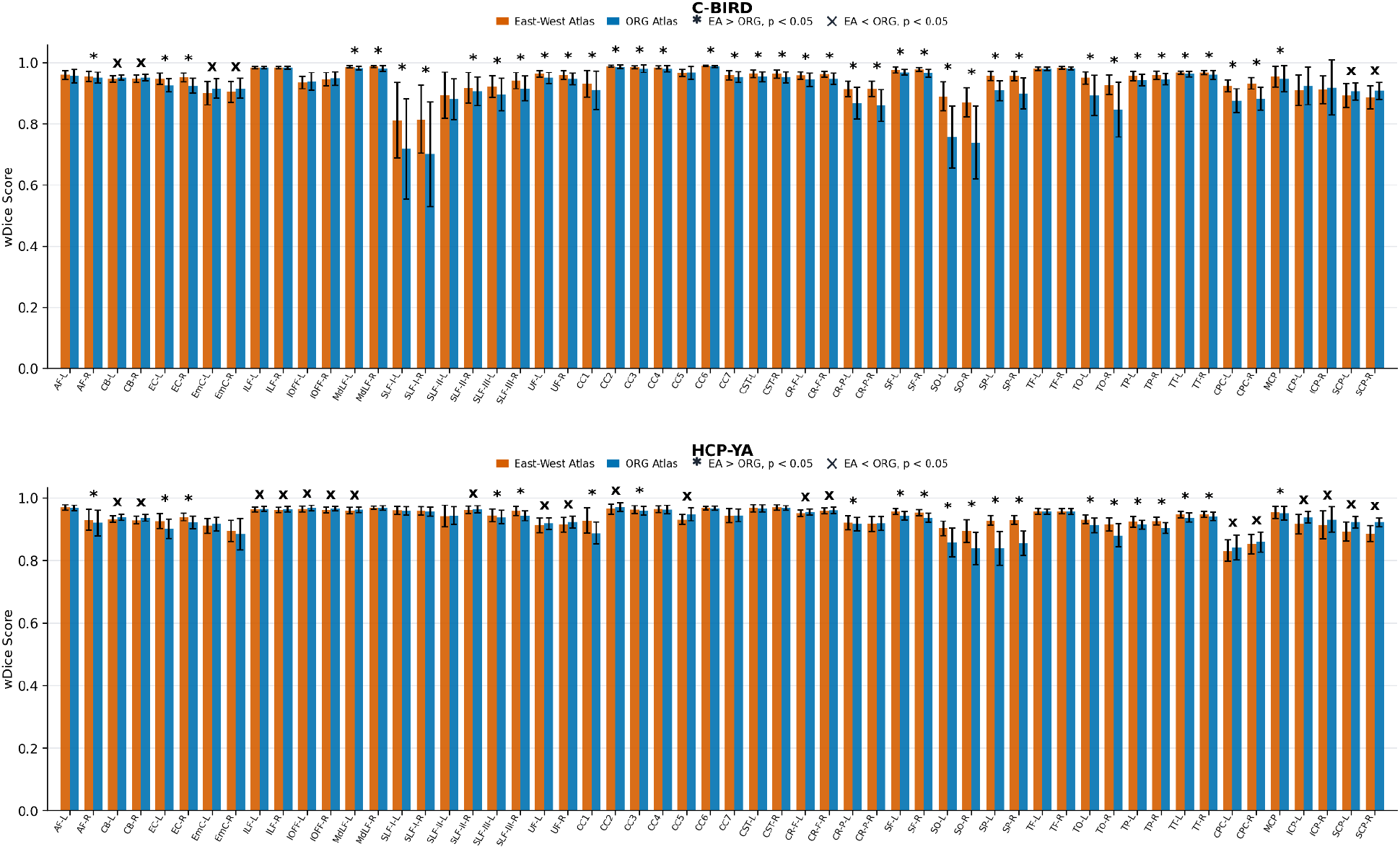
Individual-tract volumetric overlap comparison.

### 3.3 Relative difference of FA results

Figure 6 shows the overall comparison of the relative difference of mean FA between the ORG atlas and the East-West White Matter Atlas. In both the C-BIRD and HCP-YA datasets, the East-West White Matter Atlas showed lower relative difference values than the ORG atlas, indicating higher test-retest reproducibility of diffusion-derived measurements. The differences between the two atlases were statistically significant in the overall tract-level comparison for both datasets (p *<* 0.001, two-tailed paired t-test).

**Figure 6.**
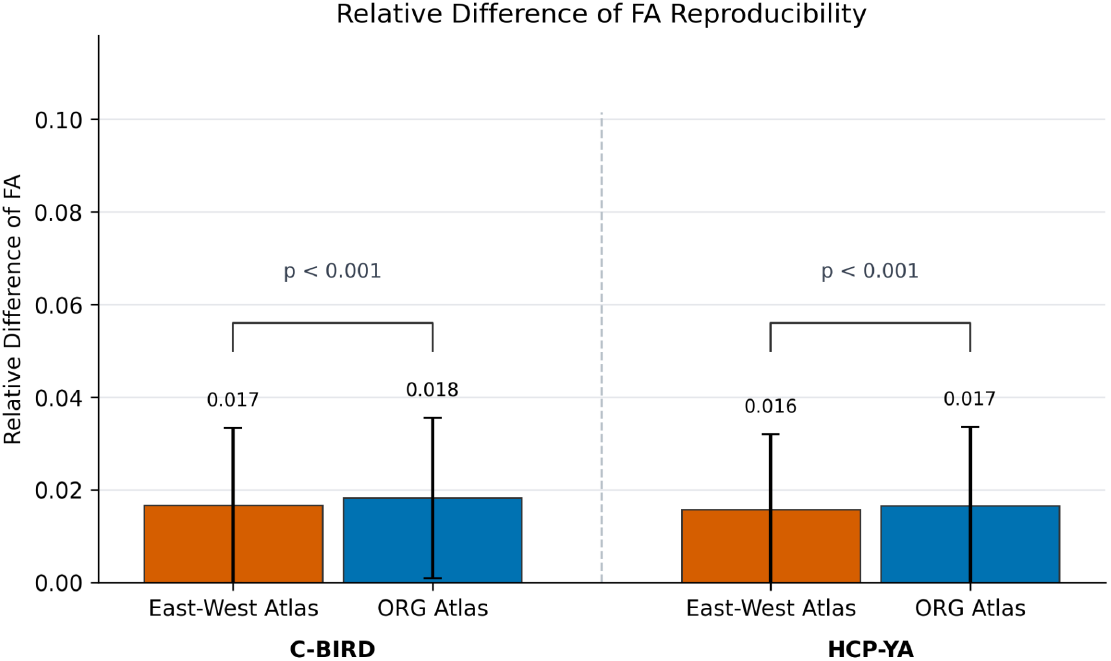
Overall comparison of the relative difference of the mean FA.

Figure 7 shows the individual-tract comparison of the relative difference of mean FA. In the C-BIRD dataset, 14 tracts showed significantly lower relative difference values using the East-West White Matter Atlas, whereas no tracts showed significantly lower values using the ORG atlas. In the HCP-YA dataset, 9 tracts showed significantly lower relative difference values using the East-West White Matter Atlas, whereas 2 tracts showed significantly lower values using the ORG atlas (p *<* 0.05, two-tailed paired t-test, FDR-corrected). These tract-level findings support the overall result that the East-West White Matter Atlas provides more stable tract-specific FA measurements across test-retest scans.

**Figure 7.**
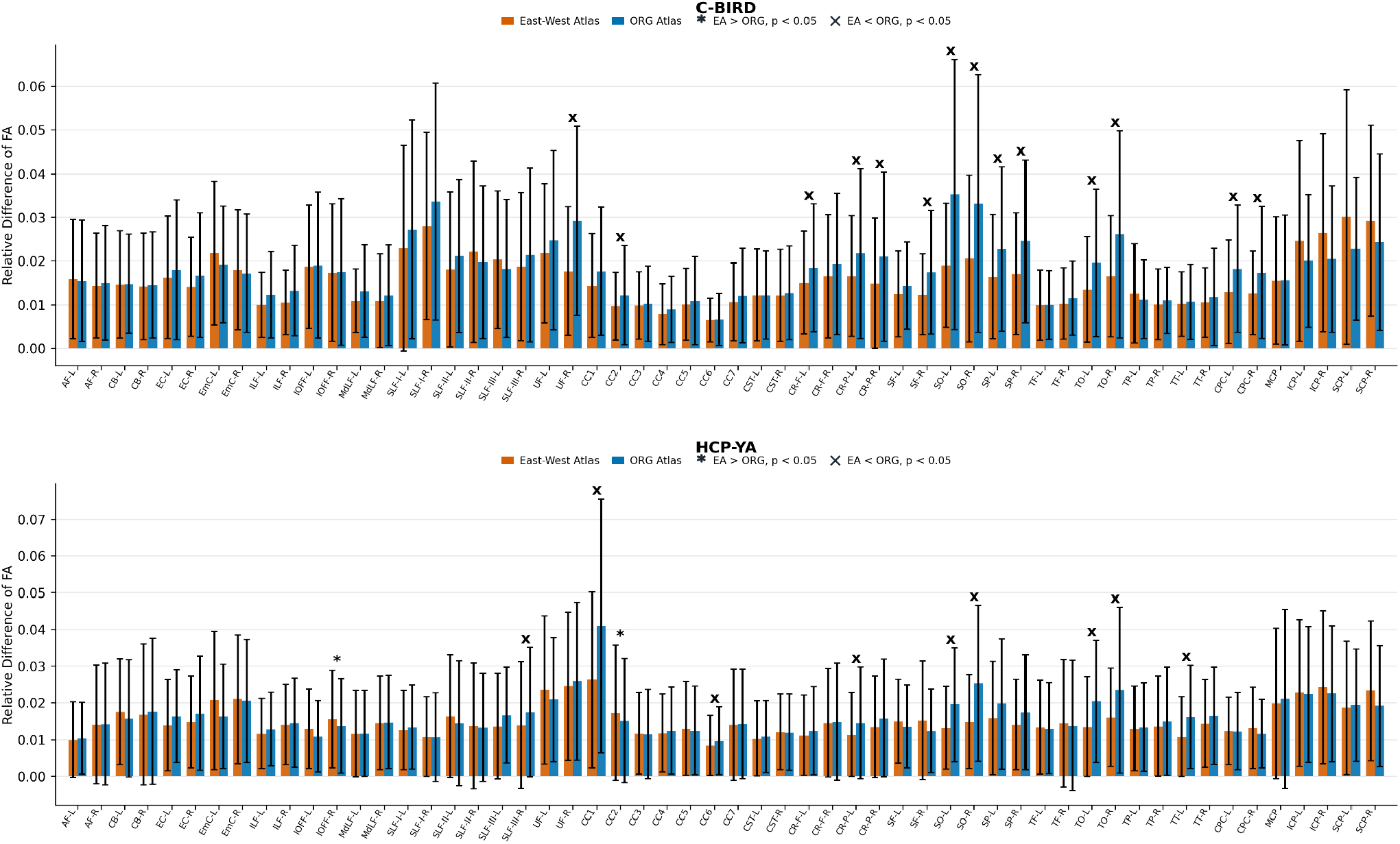
Individual-tract comparison of the relative difference of mean FA.

## 4 Discussion

In this study, we systematically evaluated the test-retest reproducibility of two clustering-based white matter atlases, the single-population ORG atlas and the cross-population East-West White Matter Atlas, using test-retest diffusion MRI datasets from two independent populations. Reproducibility was assessed from multiple complementary perspectives, including tract detection rate at the hemisphere-separated cluster and anatomical tract levels, spatial overlap measured by wDice, and diffusion-derived measurement stability quantified by the relative difference of mean FA (Zhang et al., 2019). The main findings were threefold. First, both atlases showed stable tract detection across test-retest scans at both detection levels, supporting the reliability of clustering-based atlas parcellation for identifying major white matter structures. Second, the East-West White Matter Atlas achieved higher overall wDice values than the ORG atlas in both datasets. Third, the East-West White Matter Atlas showed lower relative differences in mean FA across test-retest scans, indicating improved reproducibility of diffusion-derived tract measurements.

The detection rate results suggest that both atlases provide stable white matter structure identification across test-retest scans. At the anatomical tract level, both atlases achieved complete detection in the test and retest sessions of both datasets, indicating that both atlas frameworks can reliably identify the major deep white matter tract set in test-retest acquisitions. At the cluster-level detection rate, the East-West White Matter Atlas showed higher detection rates than the ORG atlas in the C-BIRD dataset, whereas the ORG atlas showed slightly higher detection rates in the HCP-YA dataset. This pattern is consistent with the population characteristics of the atlases and datasets. The C-BIRD dataset was acquired from an Eastern population and had lower diffusion imaging resolution and angular sampling than the HCP-YA dataset. Because the East-West White Matter Atlas was constructed using both CHCP and HCP-YA subjects, it may better capture white matter variability across Eastern and Western populations and may therefore provide broader anatomical coverage when applied to the C-BIRD dataset (Li et al., 2024). In contrast, the ORG atlas was constructed from HCP-YA subjects, making its slightly higher or comparable cluster-level detection rate in the HCP-YA dataset reasonable. These findings highlight the importance of population information in atlas construction and suggest that cross-population atlases may be particularly valuable for studies involving diverse cohorts, multi-site datasets, or populations that differ from those used to construct conventional atlases.

The wDice results further demonstrate the advantage of the East-West White Matter Atlas in terms of spatial reproducibility. The wDice measure reflects the degree of spatial overlap between corresponding parcellated tracts in test-retest scans while accounting for local fiber density (Zhang et al., 2019). In both datasets, the East-West White Matter Atlas achieved significantly higher overall wDice values than the ORG atlas, indicating more consistent spatial localization of anatomical tracts across test and retest scans. This suggests that the tract definitions in the cross-population atlas may be more robust to inter-individual anatomical variability and differences in acquisition quality. Importantly, the advantage of the East-West White Matter Atlas was observed not only in the C-BIRD dataset but also in the HCP-YA dataset. This finding indicates that incorporating an Eastern population during atlas construction did not reduce the applicability of the atlas to a Western cohort. Instead, the cross-population atlas improved spatial reproducibility across both independent datasets, supporting its generalizability for atlas-based white matter parcellation.

The relative difference of mean FA provides complementary information about the reproducibility of diffusion-derived quantitative measurements. FA is widely used in diffusion MRI studies to characterize white matter microstructural properties, and stable FA measurements are essential for reliable quantitative analyses (Boukadi et al., 2019; Laguna et al., 2020; Zhang et al., 2019). In this study, the East-West White Matter Atlas showed lower relative differences in mean FA than the ORG atlas in both datasets, indicating higher test-retest reproducibility of diffusion-derived measurements. This finding suggests that the East-West White Matter Atlas not only provides more consistent geometric tract localization but also supports more stable extraction of tract-specific diffusion metrics. Improved FA reproducibility is particularly important for longitudinal studies, multi-site studies, and clinical neuroimaging applications, where measurement variability can reduce statistical power and obscure biologically meaningful changes. Therefore, the lower FA variability observed with the East-West White Matter Atlas suggests that cross-population atlas construction may improve the reliability of downstream quantitative tractography analyses.

Although the East-West White Matter Atlas showed better overall reproducibility, the individual-tract analyses indicated that atlas performance was not uniform across all white matter tracts. The ORG atlas showed higher reproducibility for a subset of tracts, suggesting that the relative performance of different atlases may depend on tract-specific anatomical and methodological factors. Potential factors include tract size, trajectory length, curvature, anatomical complexity, and the degree of overlap with crossing-fiber regions (Maier-Hein et al., 2017; Schilling et al., 2019). Tracts located near cortical terminations or deep gray matter structures may also be more sensitive to registration variability and tractography differences. In addition, some tracts may be represented more consistently in the population used to construct one atlas than in another. These tract-specific differences indicate that atlas evaluation should not rely solely on global summary measures. Instead, both overall reproducibility and tract-level performance should be considered when selecting an atlas for a specific research or clinical application.

The present findings have important implications for reproducible quantitative tractography. By integrating data from Eastern and Western populations, the East-West White Matter Atlas may provide a more generalizable representation of white matter anatomy than a single-population atlas (Li et al., 2024; Zhang et al., 2026). Its improved performance across both the C-BIRD and HCP-YA datasets suggests that cross-population atlas construction can enhance robustness across populations and acquisition protocols without compromising performance in the original Western cohort. This is particularly relevant for large-scale neuroimaging studies, multi-center collaborations, and clinical applications, where datasets often include participants from heterogeneous populations and are acquired using different imaging protocols. More broadly, these results emphasize that white matter atlas construction should consider population diversity, especially when an atlas is intended for use across independent cohorts or imaging sites.

Several limitations should be noted. First, this study included only two young adult datasets. Although the C-BIRD and HCP-YA datasets differed in population and acquisition protocol, additional datasets covering broader age ranges, clinical populations, and more diverse imaging conditions are needed to further evaluate whether the advantages of cross-population atlas construction generalize beyond young healthy adults. Second, whole-brain tractography was reconstructed using only UKF tractography. Because tractography results can vary across reconstruction algorithms, tracking parameters, reconstruction models, and preprocessing pipelines, future studies should evaluate whether the observed atlas differences remain consistent across different tractography workflows. Third, the two datasets differed in test-retest interval and diffusion acquisition protocol, which may have contributed to differences in reproducibility measurements. Although this design allowed us to assess atlas performance under different imaging conditions, future studies with harmonized acquisition protocols would help isolate the effect of atlas construction from dataset-related variability.

## 5 Conclusion

In this study, we evaluated the test-retest reproducibility of two clustering-based white matter atlases using test-retest diffusion MRI datasets from two independent populations. Both atlases showed stable tract detection at the hemisphere-separated cluster and anatomical tract levels across test and retest scans, supporting the reliability of atlas-based white matter parcellation. Compared with the single-population ORG atlas, the cross-population East-West White Matter Atlas demonstrated higher overall spatial overlap and lower variability in mean FA measurements, indicating improved geometric and diffusion-derived reproducibility. These findings suggest that incorporating cross-population information into atlas construction can enhance the robustness and generalizability of white matter parcellation, supporting the use of the East-West White Matter Atlas for reproducible quantitative tractography analyses across diverse populations.

## Acknowledgments

This work is in part supported by the National Key R&D Program of China (No. 2023YFE0118600), the National Natural Science Foundation of China (No. 62371107), and Science and Technology Department of Sichuan Province (No. 2026YFHZ0045)

